## Supplementary material for "Dynamic Gamma Modulation of Hippocampal Place Cells Predominates Development of Theta Sequences": figure supplement

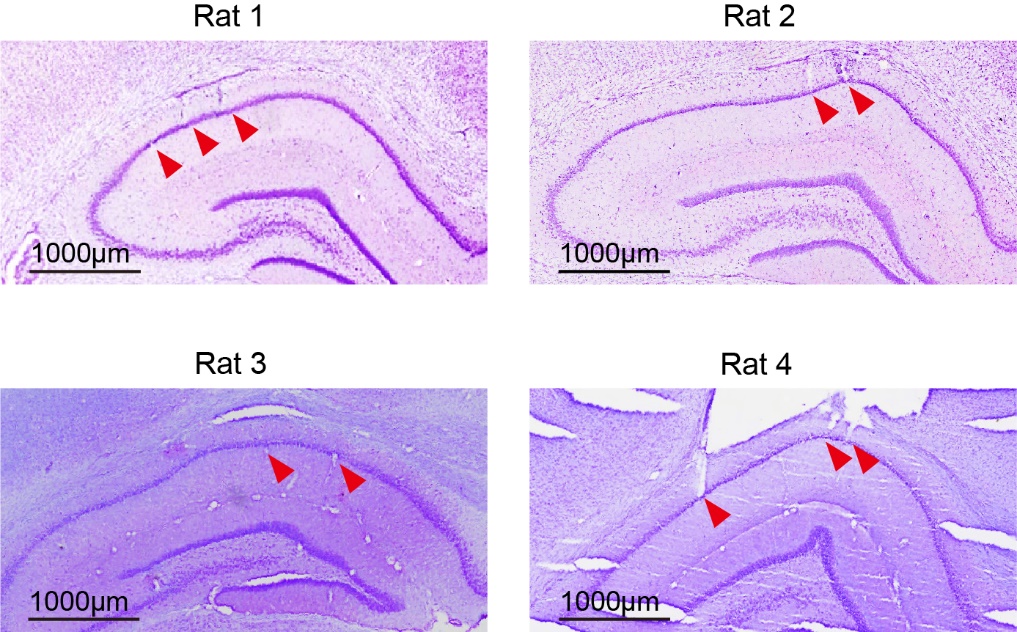

**Figure 2-figure supplement 1 | Histological verification of tetrodes.** Histologic sections showing example recording sites in CA1 of four rats. The red triangle marks the location of tetrode tip in CA1 pyramidal layer.

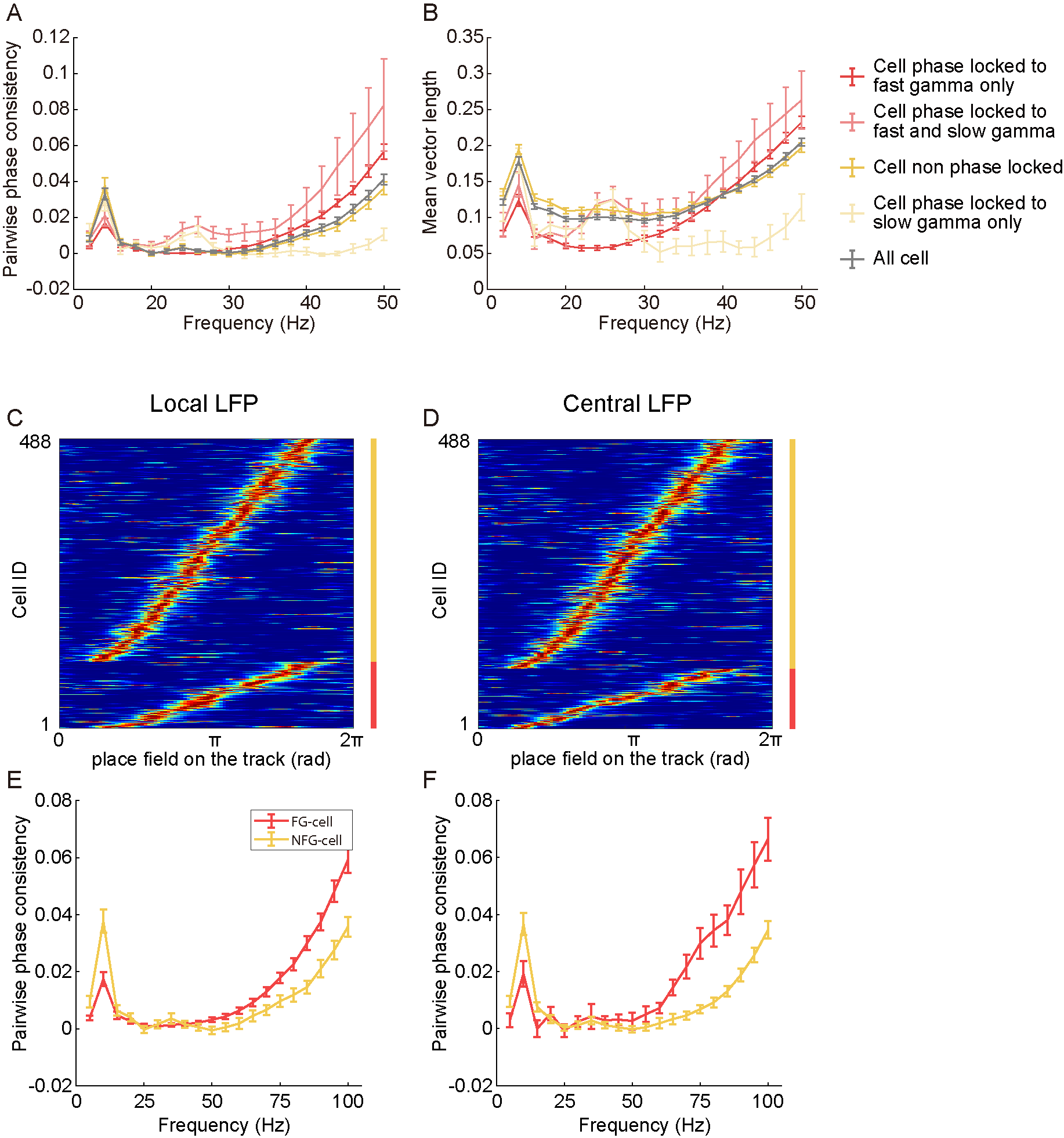

**Figure 2-figure supplement 2 | Phase locking as a function of frequency.** (A) Pairwise phase consistency (PPC) of 4 type cell of figure 2 in 5Hz steps over a range of 0-100Hz. (B) Same as (A) but for mean vector length. Legend color was same as figure 2. Grey line indicated all 488 cells. (C) and (D) the normalized turning curves of FG-cells (red) and NFG-cells (yellow) detected according to the local LFP and consistent central LFP as reference, respectively. The number of FG-cells in (C) and (D) were 113 (23.2%) and 101 (20.7%). They were sorted by the center of mass of main place field. (E) The PPC of FG- and NFG-cells in (C), modulated by all LFP frequency band (1-100Hz). (F) The PPC of FG- and NFG-cells in (D), modulated by all LFP frequency band (1-100Hz). Data are presented as mean ± SEM.

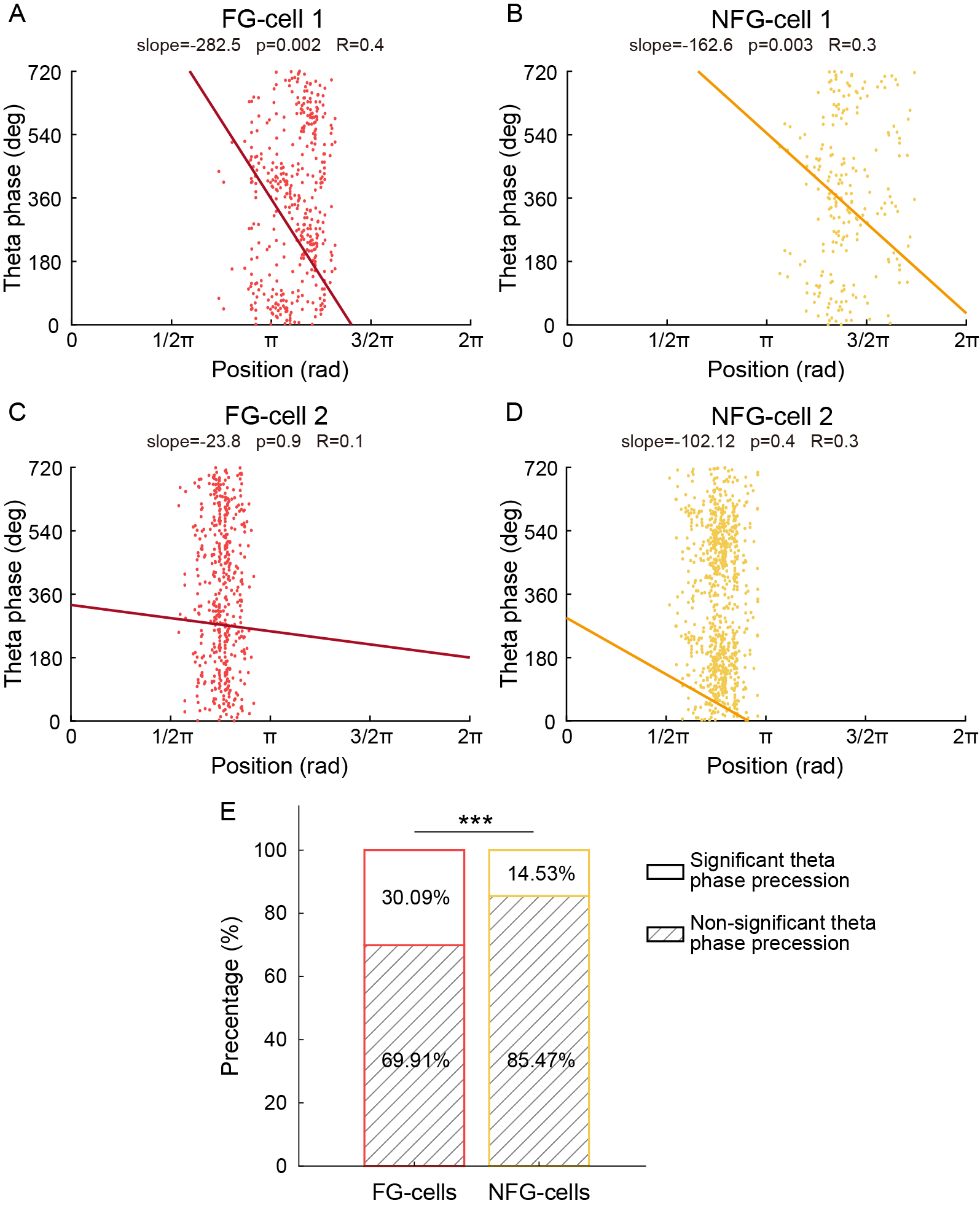

**Figure 2-figure supplement 3 | Theta phase precession of FG- and NFG-cells.** (A) An example FG-cell with significant theta phase precession. Red line shows the linear-circular regression of scatter plots. (B) An example NFG-cell with significant theta phase precession. Yellow line shows the linear-circular regression of scatter plots. (C) An example of FG-cell without significant theta phase precession. (D) An example NFG-cell without significant theta phase precession. (E) Percentage of FG- and NFG-cells with or without significant theta phase precession. ***p<0.001.

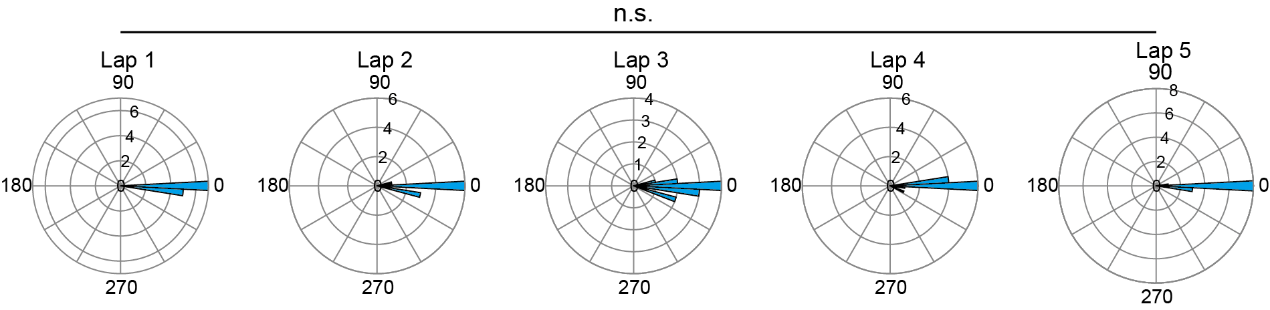

**Figure 3-figure supplement 1 | Head direction across trials.** The polar coordinates’ histogram of angle difference between instantaneous head direction and angle of tangent vector along the circular track across laps. The distribution of angle difference did not significantly change across laps.
